# Mesenchymal Stromal Cells Immunosuppress Osteoarthritis Synovial Fluid Modulated Monocytes via IL-6 and CCL2

**DOI:** 10.1101/2025.08.14.669875

**Authors:** Mozhgan Rasti, Aida Feiz Barazandeh, Kevin P. Robb, Rachel Low, Oreoluwa Kolade, Kevin Fan, Rajiv Gandhi, Sowmya Viswanathan

**Affiliations:** Schroeder Arthritis Institute, University Health Network, Toronto, Ontario, M5T 0S8, Canada; Krembil Research Institute, University Health Network, Toronto, Ontario, M5T 0S8, Canada; Institute of Biomedical Engineering, University of Toronto; Toronto, Ontario, M5S 3G9, Canada; Department of Surgery, Division of Orthopedic Surgery, University of Toronto, Toronto, Ontario, M5T 1P5, Canada; Department of Medicine, University of Toronto, Toronto, Ontario, M5S 3H2, Canada

## Abstract

**Background:** Mesenchymal stromal cell (MSC) interactions with monocytes/macrophages are central to their therapeutic effects in knee osteoarthritis (KOA); however, mechanisms of these interactions are not fully understood.

**Hypothesis:** We hypothesize that MSC soluble factors, particularly interleukin-6 (IL-6) and C-C motif chemokine ligand (CCL2) modulate monocytes in KOA environment.

**Methods:** Using healthy donor CD14^+^ monocytes exposed to KOA synovial fluid (SF) in the presence or absence of <u>m</u>arrow-derived MSC<u>(M)</u> directly or conditioned medium (CM), we evaluated cell surface markers and signaling via signal transducer and activator of transcription (STAT3), nuclear factor kappa-light-chain-enhancer of activated B (NF-κB) and c-Jun Terminal Kinase (JNK); functional responses were measured by secretion of tumor necrosis factor (TNF) and IL-1, and by phagocytosis of pHrodo Red *E-coli*.

**Results:** CD14^+^ monocytes demonstrated a mixed phenotype in KOA SF with increased CD163, CD206 and unchanged HLA-DR, CD86 marker expression. This was accompanied by activated STAT3, JNK and NF-κB signaling. TNF and IL-1 secreted levels were unchanged, but phagocytosis was impaired, indicative of a net dysfunctional repair phenotype and functionality. CD14^+^ monocytes in KOA SF were hyporesponsive to additional lipopolysaccharide re-challenge, based on TNF and IL-1 secretion. Addition of MSC(M) to KOA SF programmed CD14^+^ monocytes resolved the dysfunctional phenotype and functionality, with significant increases in CD163, CD206; significant reductions in HLA-DR and CD86 expression; this was accompanied by significantly increased activated STAT3, and decreased activated JNK and NF-κB. TNF and IL-1 secretion were also significantly reduced, and phagocytic capacity restored. Blocking IL-6, or to a lesser extent, CCL2, partially abrogated MSC(M) soluble factor effects. MSC(M) experienced apoptosis in KOA SF; however, apoptotic bodies did not fully recapitulate MSC(M) soluble factor effects.

**Conclusion:** IL-6, CCL2, other soluble factors and apoptotic bodies from MSC(M) secretome mitigate the dysfunctional effects of KOA SF on CD14^+^ monocytes resulting in immunosuppressed phenotype and functionality.

## Introduction

Osteoarthritis (OA) is a chronic, low-grade inflammatory condition that affects over 15% of the global population (1). Inflammation of the synovium lining, or synovitis (2), contributes significantly to OA pathology (3,4) with monocytes and macrophages (MΦs), being the most prevalent immune mediators in knee OA (KOA) patients’ synovial fluid (SF) (5–7). Monocytes/MΦs in the joint originate from circulating monocytes that infiltrate the joint or are residual from embryonic development (8,9); they exist in a spectrum of functional and transcriptomic states (9–13), which are driven by complex KOA multi-tissue cellular interactions, mechanical and metabolic stress (9,12), and are now acknowledged as relevant therapeutic targets for disease modification in KOA (14–16).

Mesenchymal stromal cells (MSCs) are promising cell therapies for KOA (17–22) with evidence suggesting their therapeutic potential stems from interactions with joint monocyte/MΦs (23,24). MSCs exposed to KOA SF secrete interleukin (IL)-6 and C-C motif Chemokine Ligand (CCL2) (25), which we hypothesize play key regulatory roles in the context of MSC interactions with monocyte/ MΦs in KOA.

We first describe how KOA synovial fluid (SF) causes dysfunctional re-programming of peripheral CD14^+^ monocytes, likely reflecting early changes to infiltrating monocytes in the OA joint environment. SF contains soluble mediators released from multiple joint tissues (26,27) providing an experimentally accessible proxy of KOA joint signals. Next, we show that <u>m</u>arrow-derived MSC<u>(M</u>) resolved the KOA SF induced dysfunction of CD14^+^ monocytes and these effects were mediated partially by IL-6 and CCL2 signaling.

## Results

### Late-Stage OA SF MΦs exhibit a mixed phenotypic and functional profile

SF from mid- and late-stage KOA patients **(Table S1)** showed increases in cytokines, chemokines and alarmins **(Table S2, Figs. S1A-C)**. IL-6 and S100A8/9 increased from non-diseased controls and mid-OA SF to late-OA SF **(Figs. 1A,C),** while CCL2 increased from non-diseased controls to mid-and late-OA SF **(Fig. 1B)**. Monocytes/MΦs present in late-OA SF were immunophenotyped; classical CD14^+^CD16^neg^ and intermediate CD14^+^CD16^+^ MΦs subsets were the most abundant, as before (6) **(Fig. 1D).** SF monocyte/MΦs from KOA patients expressed CD163, CD206, Human Leukocyte Antigen-DR isotype (HLA-DR) and CD80 simultaneously **(Fig. 1E)**; mixed phenotypic expression was also present in control, non-diseased SF monocyte/MΦs obtained from cadaveric donors, routinely used as controls (28), although insufficient cell numbers from cadaveric donors prevented further downstream exploration. The availability of sufficient KOA SF monocytes/MΦs allowed for further culture and re-challenge with lipopolysaccharide (LPS; late-OA SF already contains LPS (29,30)) and showed no significant phenotypic or intracellular IL-1 and TNF protein changes, suggestive of hyporesponsive KOA SF monocytes/MΦs **(Fig. 1F)**.

**Figure 1.**
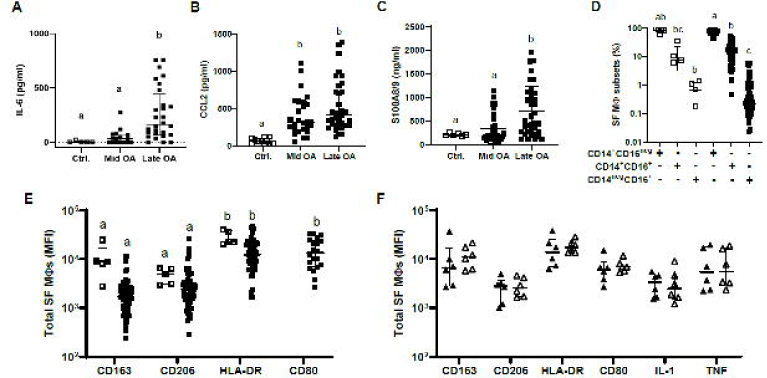
KOA synovial fluid analysis and immunophenotyping of monocytes/MΦs. **(A-C)** Median and interquartile range of IL-6, chemokine (C-C motif) ligand 2 (CCL2), and S100A8/9 concentrations in synovial fluid (SF) samples from control SF (Ctrl, non-diseased; N=6-11); mid-stage KOA SF (Mid OA; N=21-30); late-stage KOA SF (Late-OA; N=29-35). **(D)** Frequency of monocytes/MΦ subsets in the non-diseased control (□; N=5) and late-KOA SF (▪; N=64). **(E)** Mean fluorescence intensity (MFI) of surface markers on the non-diseased control (□; N=5) and KOA SF (▪; N=60) monocytes/MΦ. **(F)** Immunophenotyping, intracellular IL-1 and TNF evaluation of cultured KOA SF monocyte/MΦs (N=6) following re-challenge with (Δ) and without (▴) lipopolysaccharides (LPS). Letters indicate significant differences between groups based on the Kruskal-Wallis test, with Dunn’s multiple comparisons test.

### CD14^+^ peripheral monocytes exposed to OA SF

To understand how the OA environment modulates circulating monocytes as they infiltrate the joint, we used peripheral CD14^+^ monocytes from healthy donors or KOA patients and exposed them to late-OA SF for 1-3 days to understand these initial changes **(Fig. 2A)**. CD14^+^ monocytes from healthy or KOA patients showed similar phenotypic changes, activation of signaling pathways, secreted factors and phagocytic capacity in response to late OA SF **(Table S3)**; primary data is presented with healthy CD14^+^ monocytes; supplementary data includes both healthy and KOA CD14^+^ monocytes.

**Figure 2.**
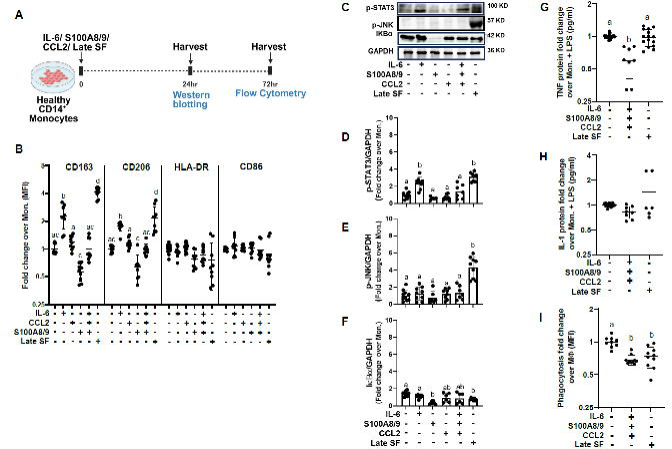
Late-OA SF reprograms CD14^+^ monocytes, partially replicated by IL-6 and CCL2 stimulation. **(A)** Experimental setup for exposure of healthy peripheral CD14 monocytes to KOA SF. **(B)** MFI of CD163, CD206, HLA-DR, and CD86 surface markers, with indicated treatments. **(C)** Representative Western blot. **(D-F)** Bar graphs illustrating phosphorylated-signal transducer and activator of transcription 3 (p-STAT3), phosphorylated-c-Jun terminal kinase (p-JNK), and nuclear factor of kappa light polypeptide gene enhancer in B-cells inhibitor alpha (IκBα) levels (normalized to glyceraldehyde-3-phosphate dehydrogenase; GAPDH). **(G, H)** TNF and IL-1 soluble factor production by CD14^+^ monocytes under the indicated treatments, normalized to control LPS stimulation. **(I)** Phagocytosis by CD14^+^ macrophages at indicated treatments. N=3 biological replicates; n≥3 technical replicates. Letters indicate significant differences based on an ordinary one-way ANOVA followed by Tukey’s multiple comparisons test.

### Late-OA SF impairs CD14^+^ monocyte functionality

Peripheral CD14^+^ monocytes from healthy donors exposed to pooled SF from late-stage KOA patients showed increased CD163, CD206 and unchanged HLA-DR and CD86 expression **(Figs. 2A,B)**. Additionally, they displayed an elevated frequency of intermediate with decreased frequency of classical monocyte subsets **(Figs. S2A,B)**, like frequency changes between SF vs. circulating monocytes/MΦs in KOA patients (6). Phenotypic changes of KOA patient vs. healthy donor sourced CD14^+^ monocytes were broadly comparable **(Table S3)**.

Next, we examined the effects of IL-6, S100A8/9, an endogenous TLR ligand, and CCL2 individually and in combination to mimic the observed effects of late-OA SF on CD14^+^ peripheral monocytes. IL-6, CCL2 and S100A8/9 were elevated in late-OA SF **(Figs. 1A-C)**; IL-6 and CCL2 are of interest given their roles in KOA but also their paradoxical expression by MSCs exposed to KOA SF (25). Physiological levels of IL-6, CCL2 and S100A8/9 based on KOA SF levels were used for experiments, although hyperphysiological levels of IL-6 but not CCL2 elicited additional phenotypic changes **(Figs. S2C,D)**. We confirmed CD14^+^ monocytes from healthy and KOA patients express membrane bound IL-6R (mIL-6R), which increases with physiological IL-6 or late-OA SF stimulation, and also express C-C Receptor 2 (CCR2), unchanged by physiological or late-OA SF stimulation **(Figs. S2E,F**). Specifically, IL-6, S100A8/9 and CCL2 individually mimicked some, but not all aspects of late-OA SF **(Fig. 2B, Table S4)**. The combination of all three factors also failed to fully mimic phenotypic changes in healthy CD14^+^ monocytes **(Fig. 2B, Table S4)**, suggestive of other factors in KOA SF that work together with IL-6, CCL2 and S100A8/9.

To understand CD14^+^ phenotypic changes to late-OA SF, we examined kinase activation using a phospho-kinase array **(Fig. S3A)**. Significant activation of signal transducer and activator of transcription (STAT3) and c-Jun Terminal Kinase (JNK), amongst multiple kinases evaluated, was evident **(Figs. S3A-E)**. Peripheral CD14^+^ monocytes from healthy **(Figs. 2C-F)** and KOA patients **(Figs. S5B-E)** showed increased p-STAT3 and p-JNK and reductions in negative canonical regulator of bound nuclear factor of kappa light polypeptide gene enhancer in B-cells (NF-κB), inhibitor alpha (IκBα) (31) upon late-OA SF stimulation. Recombinant IL-6, CCL2, S100A8/9 or their combinations did not fully mimic the effects of late-OA SF on healthy CD14^+^ monocytes **(Figs. 2C-F, Table S4).**

We next evaluated functionality of CD14^+^ monocytes exposed to late-OA SF and observed no changes in tumor necrosis factor (TNF) or IL-1 secretion **(Figs. 2G,H)**. Recombinant IL-6 however, reduced TNF and IL-1 secretion, while recombinant CCL2 only reduced IL-1 secretion **(Figs. S2G,H)**; the combination of all three proteins reduced TNF secretion **(Figs. 2G,H, Table S4)**, suggestive of other co-factors in KOA SF that masked the observed effects of these three factors.

Despite limited effects on cytokine secretion, late-OA SF or the three factors in combination significantly impaired phagocytotic capacity of CD14^+^ monocytes **(Fig. 2I)**. Taken together, CD14^+^ monocytes displayed limited pro-inflammatory changes but exhibited reduced functionality to late OA SF.

To further test the limited response of CD14^+^ monocytes to KOA SF, we re-challenged healthy and KOA CD14^+^ monocytes with LPS and observed mixed shifts in their phenotypes and no increases in TNF or IL-1 secretion, suggestive of hyporesponsive functional states (**Fig. S4**). Interestingly, peripheral CD14^+^ monocytes mimicked the limited responsiveness of SF MΦs to LPS-rechallenge **(Fig. 1F)**, suggestive of desensitization to the late-OA SF chronic, low-grade inflammatory milieu.

### Effects of IL-6, CCL2 or TLR inhibition on CD14^+^ monocytes

Given partial effects of IL-6, CCL2 and S100A8/9 on CD14^+^ monocytes, we evaluated inhibition, using neutralizing antibodies against IL-6, CCL2, or a TLR4 inhibitor, TAK242 (32), in the presence of late-OA SF **(Fig. 3A)**. IL-6 neutralizing antibody abrogated late-OA SF mediated changes on CD163, while CCL2 neutralizing antibody reduced CD206; TAK242 did not produce any phenotypic effects **(Figs. 3B, S5A)**. IL-6 and CCL2 neutralizing antibody effects were largely mediated by suppressing p-STAT3; IL-6 but not CCL2 neutralizing antibody also reduced IκBα and increased p-JNK levels **(Figs. 3C-F)**. Surprisingly, TLR antagonism via TAK242 did not produce any effects on JNK activation but increased IκBα **(Figs. 3C,E, F)**. IL-6 antibody suppression had similar effects on KOA sourced CD14^+^ monocytes **(Figs. S5B-E)**.

**Figure 3.**
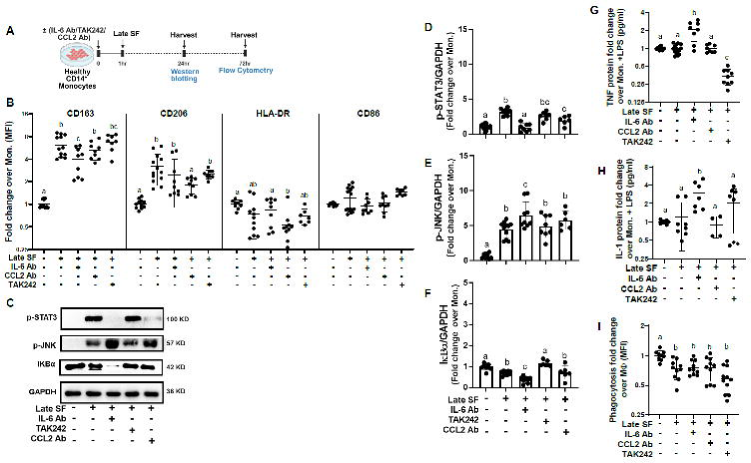
Blocking IL-6, CCL2, S100A8/9 in late-OA SF programed CD14^+^ monocytes. **(A)** Experimental timeline for blocking healthy CD14 monocytes. **(B)** MFI of CD163, CD206, HLA-DR, and CD86 surface markers, on CD14 monocytes at the indicated treatments. **(C)** Representative Western blot. **(D-F)** Bar graphs of p-STAT3, p-JNK, and IκBα (normalized to GAPDH). **(G, H)** TNF and IL-1 soluble factor production at indicated treatments, normalized to control LPS stimulation. **(I)** Phagocytosis by CD14^+^ macrophages at the indicated treatments. N=3 biological replicates; n=3 technical replicates. Letters indicate significant differences between groups based on an ordinary one-way ANOVA followed by Tukey’s multiple comparisons test.

Addition of IL-6 but not CCL2 neutralizing antibody to late-OA SF increased TNF and IL-1 secretion **(Figs. 3G,H)**. TAK242 inhibitor decreased TNF but not IL-1 **(Figs. 3G-H)**, confirming roles for TLR4 pathway in TNF but not in IL-1 production in late-OA SF. None of the blockers individually reversed the functional phagocytotic impairment induced by late-OA SF **(Fig. 3I)**.

Of all the inhibitors, IL-6 neutralizing antibody had multiple effects on CD14^+^ monocytes. We thus examined IL-6 signaling further. Canonical IL-6 signaling occurs through mIL-6 receptor while trans-signaling occurs through soluble IL-6 receptor (sIL-6R) and gp130 **(Fig. S6A)**. IL-6 levels increased from mid- to late-stage OA **(Fig. 1A)** but sIL-6R levels decreased **(Fig. S6B)**, arguing against IL-6 trans-signaling in late-OA SF. Physiological levels of IL-6 with or without physiological sIL-6R did not impact phenotype in healthy CD14^+^ monocytes, confirming absence of competitive trans-signaling **(Figs. S6C-F)**. This was replicated by the addition of soluble gp130 (sgp130) in the presence of late-

OA SF **(Figs. S6C-F)**. Additionally, mIL-6R expression increased on CD14^+^ monocytes exposed to late-OA SF **(Fig. S2E)**, confirming dominance of classical IL-6 signaling. There was a complete absence of detectable IL-6-sIL-6R complexes in both mid- and late-OA SF (undetectable in 91 of 96 samples), readily found in inflammatory arthritic controls **(Fig. S6G)**. Absence of competitive trans-signaling was also confirmed in KOA sourced CD14^+^ monocytes **(Figs. S6C-F)**.

Taken together, late-OA SF exposure activated STAT3 and JNK; reduced IκBα; did not impact TNF or IL-1 secretion but impaired phagocytosis capacity, largely mediated through IL-6 classical signaling suggestive of a hyporesponsive CD14^+^ monocyte functional state.

### MSC(M) countered late-OA SF dysfunctional effects on CD14^+^ monocytes

Healthy and KOA CD14^+^ monocytes in late-OA SF, co-cultured directly with MSC(M) showed significantly increased CD163, CD206 and significantly reduced HLA-DR, CD86 **(Figs. 4A,B, S7A,B)**.

**Figure 4.**
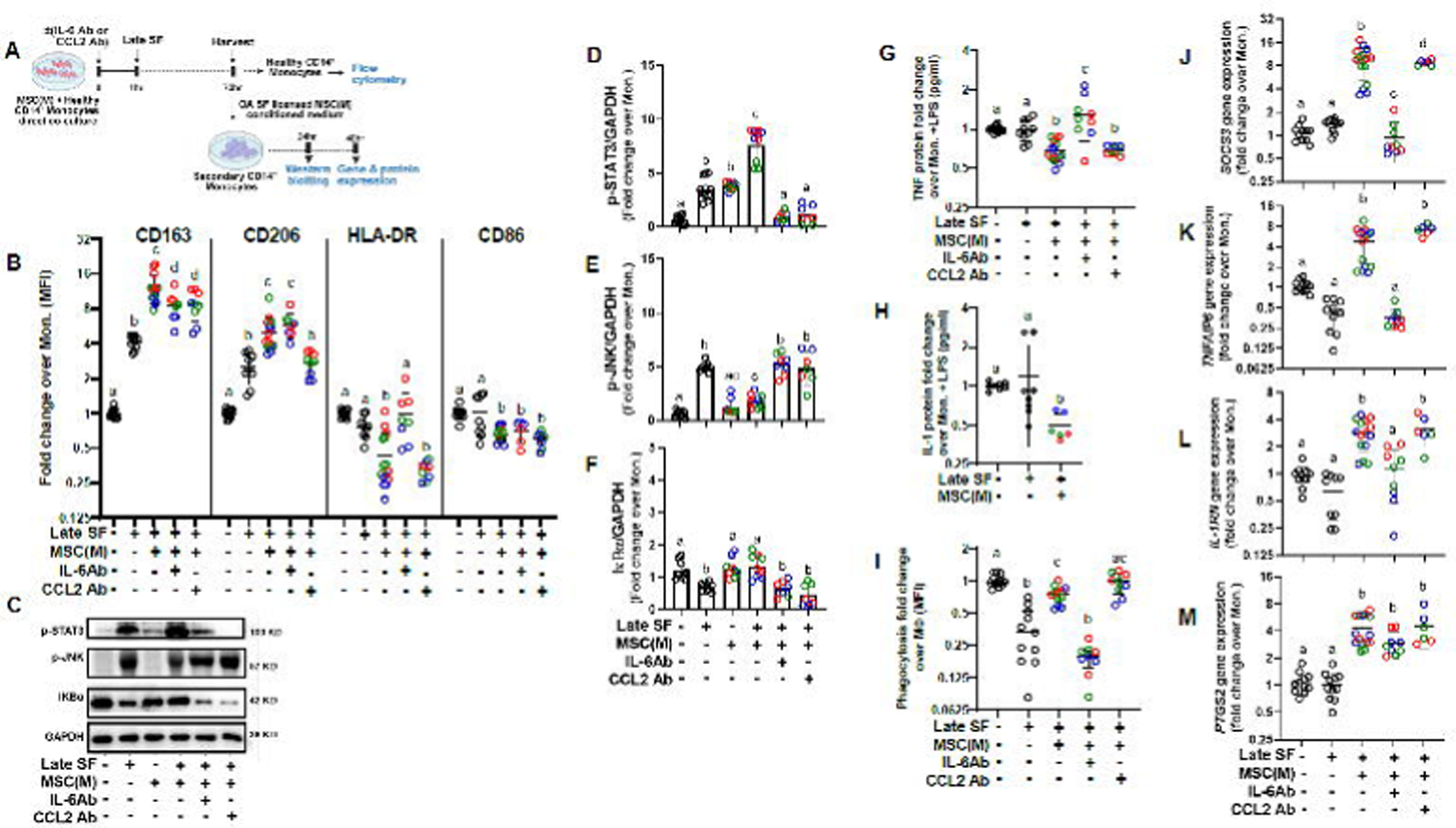
MSC(M) effects on CD14^+^ monocytes. **(A)** Experimental schematic workflow. **(B)** MFI of CD163, CD206, HLA-DR, and CD86 surface markers on healthy peripheral CD14 monocytes directly co-cultured with MSC(M), with or without late-stage OA SF. **(C)** Representative Western blot. **(D-F)** Bar graphs of p-STAT3, p-JNK, and κBα levels (normalized to GAPDH). **(G, H)** Normalized TNF and IL-1 soluble factors production by CD14^+^ monocytes in indicated treatments, with LPS spike. **(I)** Phagocytosis, at indicated treatments. **(J-M)** Expression of the suppressor of cytokine signaling 3 (*SOCS3*), TNF-alpha induced protein 6 (*TNFAIP6)*, IL-1 receptor antagonist (*IL-1RN),* and prostaglandin-endoperoxide synthase 2 (*PTGS2)* genes. MSC(M) donors are represented by different colors (blue, green, red). N=3 biological replicates; n=3 technical replicates. Letters indicate significant differences between groups based on an ordinary one-way ANOVA followed by Tukey’s multiple comparisons test.

IL-6 neutralizing antibody partially reversed MSC(M) mediated effects on CD163 and HLA-DR while CCL2 antibody reversed MSC(M) effects on CD163 and CD206 **(Figs. 4B, S7B, Table S5)**. Given that S100A8/9 did not antagonize MSC(M) phenotypic effects on CD14^+^ monocytes, **(Fig. S7C)**, it was not further investigated.

To test if these effects were mediated by MSC(M) soluble factors, secondary cultures with healthy or KOA CD14^+^ monocytes were treated with conditioned medium (CM) from primary direct co-cultures of corresponding CD14^+^ monocytes and MSC(M), in late-OA SF **(Figs. 4A, S7A)**. CD14^+^ monocytes in late-OA SF licensed MSC(M) CM were larger and more spindle-shaped **(Fig. S8A)**. Licensed MSC(M) CM significantly increased activated STAT3 and IκBα and reduced activated JNK levels in healthy and KOA CD14^+^ monocytes, relative to late-OA SF **(Figs. 4C-F, S7D-G, Table S5)**. TNF and IL-1 were attenuated by licensed MSC(M) CM and phagocytosis functionality was rescued. Both IL-6 and CCL2 neutralizing antibodies reversed MSC(M) CM effects on activated STAT3, JNK and IκBα levels **(Figs. 4C-F, S7D-G, Table S5)**. Blocking IL-6 but not CCL2 also abolished MSC(M) effects on TNF secretion and phagocytotic functionality **(Figs. 4G,I, S7H,J, Table S5)**.

Downstream effectors, including suppressor of cytokine signaling 3 (*SOCS3),* TNF-alpha Induced Protein 6 (*TNFAIP6*), IL-1 Receptor Antagonist (*IL1RN*), prostaglandin-endoperoxide synthase 2 (*PTGS2*) were significantly upregulated by licensed MSC(M) CM, and downregulated by IL-6 antibody (except *PTGS2*); CCL2 antibody downregulated *SOCS3*, but not other genes **(Figs. 4J-M, S7K-N)**.

18-21% of MSC(M) in late-OA SF demonstrated significant early apoptosis, in a donor-dependent manner, which was partially blocked by IL-6 antibody; S100A8/9 was also checked and did not induce MSC(M) apoptosis **(Fig. S8B)**. Apoptotic bodies (ABs) only slightly mimicked MSC(M) CM effects on CD14^+^ monocyte TNF production, emphasizing dominant effects of MSC(M) soluble factors **(Fig. S8C).**

## Discussion

We evaluated KOA and healthy CD14^+^ peripheral monocytes exposed to complex mixture of factors in late-OA SF to understand the basal profile of circulating monocytes in OA SF, and the rescue effects of MSC(M). Insights from our study thus provides mechanistic basis for clinically reported MSC(M) analgesic and functional improvement effects in KOA patients (19,22) through the lens of their known interactions with monocytes/MΦs (22–24).

We used peripheral CD14^+^ monocytes exposed to late-OA SF to study the effects of OA environment on monocytes as they infiltrate the joint. CD14^+^ peripheral monocytes are widely used to study OA monocytes (33,34). Importantly, circulating monocytes are precursors of recruited macrophages during KOA as our data (*unpublished*) and others (9,35) show. Thus, our study of peripheral monocytes exposed to OA environment informs early changes in precursors to synovial macrophages and is an important aspect of understanding macrophage biology in KOA.

Late-OA SF contains numerous factors including IL-6, CCL2 alarmins, endotoxins (6,30,36); pro-inflammatory cytokines (5,37); complement activators (38) and ECM fragments (39). These factors contribute to a dysfunctional state in CD14^+^ peripheral monocytes as evidenced by sundry expression of cell surface markers; simultaneous activation of STAT3, JNK and NF-κB; non-significant changes in TNF and IL-1 production but significant functional phagocytosis impairment.

In chronic inflammatory conditions, persistent exposure is expected to reshape monocyte functionality, for example, through increased inflammation via trained immunity as seen in obesity (40,41), atherosclerosis (42,43) and systemic lupus erythematosus (44,45); monocytes mount exacerbated responses to pro-inflammatory re-stimuli. However, in KOA environment, re-challenge of CD14^+^ monocytes did not result in significant TNF or IL-1 production, suggestive of a hyporesponsive rather than trained immunity state concordant with our data on SF monocytes/MΦs, which also appeared hyporesponsive to LPS re-challenges.

We examined the individual roles of IL-6, CCL2, and S100A8/9, known to be elevated in KOA SF, to see if these factors mimic the late-OA SF induced dysfunctional state in CD14^+^ monocytes. IL-6 recapitulated some effects of late-OA SF effects; blockade of IL-6 also abrogated many of late-OA SF effects, speaking to synergistic interplay between endogenous IL-6 and other factors in late-OA SF. IL-6 has paradoxical roles; it correlates with increased severity, pain and worsening function in KOA patients (46–49). IL-6 knockout mice develop more severe spontaneous OA (50), but in experimentally induced OA, there is increased cartilage damage (51), supportive of context-dependent IL-6 signaling in OA. IL-6 induces classical signaling (52) in limited cell types including monocytes/MΦs, which express mIL-6R (53–55) resulting in anti-inflammatory effects (56). Signaling through sIL-6R, generated by proteolytic cleavage of mIL-6R or through transcription of alternatively spliced IL-6R mRNA (57,58) results in trans-signaling (59), which is broader given the ubiquitous expression of gp130; trans-signaling results in stronger activation of intracellular signaling (60) and is reported to be pro-inflammatory (56). Our data supports IL-6 classical signaling to CD14^+^ monocytes, in late-OA SF.

We also tested CCL2, a chemokine important in the egress of bone marrow monocytes (61), including to the joint (62). CCL2 blockade had a limited impact on CD14^+^ functional states, consistent with limited effects observed in OA injury in CCR2 null mice (63–65). The use of physiological doses, lower than that typically used (66) may have tempered our results, Similarly, alarmin-based TLR4 activation showed only partial recapitulation of late-OA SF effects via IκBα but not JNK activation, suggestive of canonical TLR4 signaling (67) in late-OA SF through NF-κB but not JNK pathways with possible inflammasome involvement as indicated by TLR4-independent expression of IL-1 (68,69).

The combination of IL-6, CCL2, S100A8/9 did not equal the sum of its parts, nor did it fully mimic late-OA SF effects on CD14^+^ monocytes; interestingly it did impair CD14^+^ phagocytic capacity, aligned with late-OA SF effects. This contrasts with increased phagocytosis reported with chronic S100A9 homodimer stimulation during monocytes-to-macrophage differentiation (69); in our triple combination, S100A8/9 heterodimers with IL-6 and CCL2 reduced phagocytosis, relative to untreated controls, likely due to differences in experimental set-ups. Strikingly, our system and van Kooten’s (69) showed that despite chronic exposure to alarmins and other pro-inflammatory factors in late-OA SF, CD14^+^ monocyte responses are either muted or pro-resolving.

Healthy and OA CD14^+^ monocytes exposed directly to MSC(M) in late-OA SF were demonstrably skewed towards a more inflammation resolving phenotype, had decreased TNF and IL-1 secretion and improved phagocytotic functionality. IL-6 antibody abrogated multiple, but not all MSC(M) effects. MSC(M) based IL-6 effects were mediated through the JNK/STAT3 axis by reducing activated JNK and increasing activated STAT3 and IκBα. IL-6 classical signaling resulted in both increased STAT3 phosphorylation and *SOCS3*, a known modulator of IL-6 and STAT3 signaling (70), mirroring SOCS3-dependent MSC mitigation of LPS-activation of alveolar macrophages (71). CCL2 blockade of MSC(M) mimicked IL-6 blockade in terms of effects on JNK/STAT3 and IκBα, but did not impact TNF secretion or phagocytosis, unlike IL-6 neutralization.

Given that 18-21% of MSC(M) exposed to late-OA SF were undergoing early apoptosis, we cannot rule out efferocytosis of MSC(M) by CD14^+^ monocytes under direct co-culture as previously reported (23,72,73). However, late-OA SF licensed MSC(M) CM recapitulated functional, signaling and transcriptomic effects, speaking to the dominant effects of MSC(M) soluble factors. Additionally, depleting apoptotic bodies from MSC(M) CM only slightly impacted MSC(M) effects, confirming limited apoptotic body effects. Our data thus supports soluble mediators in the non-apoptotic vesicular compartment, with IL-6 and CCL2, as prime mediators of MSC(M) effects.

This study has a few limitations. We conducted detailed analyses on the individual and combined effects of IL-6, CCL2, S100A8/9 on CD14^+^ monocytes, but were not able to replicate the full effects of late-OA SF. OA SF is a complex environment with multiple fluctuating factors, which is not fully recapitulated in our three factor system. Nonetheless, our data provides novel insights into initial responses of circulating monocytes to an OA environment and shows that even with relatively short exposure periods (24-72h), there is an altered dysfunctional repair state. Addition of MSCs restored CD14^+^ monocytes to more functional reparative states, via IL-6 classical and CCL2 signaling, and by mitigating some S100A8/9 effects; these however did not fully reproduce the extent of MSC(M) immunosuppressive effects. This is highly suggestive of other MSC soluble factors that were responsible for observed effects. Next steps include detailed proteomic analyses of the SF licensed MSC(M) soluble fractions. Donor effects at the level of both CD14^+^ monocytes and MSCs were incorporated to account for inter-donor variability, while response magnitude varied; treatment-associated effects remained consistent across donors.

## Key Takeaways

- KOA SF contains monocytes/MΦs that exhibit a mixed phenotype and are hyporesponsive to additional LPS stimulation
- CD14^+^ monocytes from healthy or KOA patients, exposed to KOA SF similarly exhibited a dysfunctional repair state.
- IL-6, CCL2 or S100A8/9 alone and to a limited extent in combination mimicked some, but not all aspects of KOA SF induced CD14^+^ monocytes dysfunctionality
- MSC(M), through soluble factors resolved the altered phenotype and functionality of CD14^+^ monocytes in a KOA environment

## Materials and methods

### SF sample collection

Up to 5 ml of SF were collected from KOA patients (N=139, REB 14-7483) or non-OA screened cadaveric donors ((74), N=11, REB 24-566). KOA patients **(Table S1)** met the American College of Rheumatology criteria for symptomatic KOA; psoriatic (N=2) and rheumatoid arthritis SF (N=1) were collected under REB 21-5591 and REB14-7483, respectively. All samples were collected with informed consent.

### SF MΦs immunophenotyping

SF samples exhibiting visible blood contamination were excluded; SF monocytes/MΦs (64 late OA, 5 non-OA) were immunolabeled as before (6) **(Table S6)**. Isolated SF cells (N=6) were cultured and challenged with LPS (10 ng/mL) for 4h.

### Isolation and culture CD14^+^ monocytes and MSC(M)

Human peripheral blood mononuclear cells (PBMCs) were isolated from whole blood of healthy donors or late-stage KOA patients (REB14-7483), obtained with informed consent, according to manufacturer’s protocol. CD14□ monocytes were purified via magnetic sorting using CD14^+^ MicroBeads (Miltenyi Biotec), following the manufacturer’s protocol (75) and cryopreserved until use. CD14□ monocytes were cultured in RPMI1640 medium supplemented with 10% fetal bovine serum, 1% sodium pyruvate, and 1% penicillin-streptomycin (Gibco). CD14 monocytes (2.5 × 10□ cells/400 μL) were cultured in 24-well plates and treated with either i) human recombinant IL-6 (PeproTech, 200-06) at 0.5 ng/ml; ii) S100A8/9 (R&D Systems, B226-S8) at 1 μg/mL; iii) CCL2 (PeproTech, 300-04) at 1.5 ng/ml; iv) sIL-6R (Cedar Lane, CLCYT286) at 10 ng/mL; v) sgp130 (R&D Systems, 228-GP) at 10 ng/mL; vi) 30% (v/v%) pooled late-stage OA SF, collected from eight patients. For inhibitors, CD14^+^ monocytes were pre-treated for 1 hour with either i) IL-6 neutralizing antibody (2.5 μg/mL; R&D Systems, MAB206); ii) CCL2 neutralizing antibody (1 μg/mL; R&D Systems, MAB679) or iii) TAK-242 (1 μM/mL; Millipore-Sigma, 5.08336), followed by pooled late-state OA SF for 72 hours. Three independent biological donors with three technical replicates were used for all conditions.

MSC(M) were isolated from healthy bone <u>m</u>arrow aspirates (REB 16-5493), obtained with informed consent, expanded as before (76) and used at passage 4, with characterization as before (25). Three biological donors were used **(Table S1)**. CD14 monocytes were co-cultured directly with MSC(M) at a 5:1 ratio at indicated conditions for 72 hours. MSC(M) were licensed with pooled late-stage OA SF, with or without pre-treatment (for 1h) with IL-6 or CCL2 neutralizing antibodies. CM, termed as late-OA SF licensed MSC(M) CM, was collected for subsequent use in secondary CD14□ monocyte cultures.

### Soluble factors

Clear SF samples after centrifugation (12,000xg for 15 min) or CM were stored at −80°C until analysis. IL-6, CCL2, S100A8/9, IL-6: sIL-6R, sIL-6R, IL-1 and TNF were detected as per ELISA kit instructions **(Table S7)**. Monocytes were spiked with LPS (*E. coli* origin, 2.5 ng/mL; Millipore-Sigma) for 4 hours before collecting the CM at 48h.

### MSC(M) CM Apoptotic Body (AB) Isolation

Licensed MSC(M) CM was collected after 48h in low-particulate RoosterCollect medium (RoosterBio, SKU# M2001), centrifuged at 1,000g for 10 min, then 2,000g for 20 min to remove cell debris. ABs were isolated by ultracentrifugation at 16,000g for 30 min and resuspended in cold PBS for use. AB-free fraction was collected from the supernatant of the same spin step. All centrifugation steps were conducted at 4□.

### CD14^+^ monocytes flow cytometry

CD14^+^ monocytes were immunolabeled with indicated antibodies **(Table S6)**. MFI was quantified using FlowJo analysis software v.10.10. and normalized to the donor-matched average MFI value of the corresponding marker in untreated CD14^+^ monocytes.

### Western blot

CD14^+^ monocytes were lysed, and protein concentration was measured by a Pierce BCA kit (ThermoFisher Scientific). 15μg of cell lysates were subjected to electrophoresis and membranes were immunoblotted with primary antibodies **(Table S8);** signals were detected with a Bio-Rad chemiluminescent imaging system. Images were quantified with ImageJ analysis software. Phosphorylated kinases were normalized to glyceraldehyde-3-phosphate dehydrogenase (GAPDH) as a loading control or corresponding total protein.

### Gene expression

Total RNA was extracted with RNeasy kit (Qiagen, 73404); cDNA was synthesized using SuperScript IV VILO master mix (ThermoFisher Scientific) per the manufacturer’s instructions. The primer sequences are listed **(Table S9)**. Levels of mRNA were normalized to three housekeeping genes GAPDH, beta-2-microglobulin (B2M) and β-Actin using the 2^-ΔΔCt^ method.

### Phagocytosis

CD14 monocytes were differentiated into macrophages over 5 days with 20 ng/mL M-CSF (PeproTech). 1×10^5^ macrophages were treated for an additional 3 days with M-CSF, with or without either late-stage OA SF, or MSC(M) CM, in the presence or absence of IL-6 or CCL2 neutralizing antibodies. Phagocytosis of pH Rodo™ Deep Red *E. coli* BioParticles® Conjugate (ThermoFisher Scientific, P35360) assessed as per manufacturer’s protocol.

### Proteome profiler array

The phosphorylation of protein kinases in healthy peripheral CD14⁺ monocytes treated with late-OA SF for 1 hour was assessed using the Human Phospho-Kinase Array Kit (R&D Systems; ARY003C), according to the manufacturer’s instructions. Signal intensities, representing the relative levels of phosphorylated proteins bound to each pair of duplicate spots, were quantified with ImageJ.

### Apoptosis Assay

MSC(M) were treated with pooled late-OA SF, and the percentage of early apoptotic cells was subsequently assessed by Annexin V-FITC Apoptosis Detection Kit with 7-AAD (Biolegend, Cat# 640922), according to the manufacturer’s instructions.

### Statistical analysis

All analyses were performed using Prism (GraphPad). Normality and log normality tests were conducted prior to each analysis. Parametric data with equal variance were analyzed using ordinary one-way ANOVA with Tukey’s multiple comparisons test. For non-parametric data, verified by residual analysis, Kruskal-Wallis test with Dunn’s multiple comparisons tests were applied. Data is shown as mean ± SD of N=3 biological replicates; n=3 technical replicates, however all statistics were only performed on biological replicates. Letters indicate significant differences (p< 0.05) between groups, with the shared letters indicating no significant difference.

Supplemental material includes Fig. S1-S8 and Tables S1-S9.

## Supporting information

Graphical Abstract

## Acknowledgements

This work is supported by the Arthritis Society (YIO-15-321); CIHR (PJT-166089) to SV and Schroeder Arthritis Institute via the Toronto General and Western Hospital Foundation (University Health Network). We thank the UHN Biobank, led by Dr. M. Kapoor and team, specifically Kim Perry, Christina Ward, and Luis Montoya for sample management, consent, and data collection; Dr. V. Chandran for psoriatic and rheumatoid arthritis samples; Dr. R. Krawetz for cadaveric samples Schematics developed using BioRender.com.

## Author Contributions

Methodology, investigation, formal analysis and visualization, MR. Methodology, ABF. Writing original draft, formal analysis and visualization, KR. Visualization and editing, LR. Methodology, KF. Formal analysis, OK. Resources, fund acquisition, RG. Conceptualization, writing, original draft and editing, supervision, project administration, fund acquisition, SV.

