## supplemental files for "Mesenchymal Stromal Cells Immunosuppress Osteoarthritis Synovial Fluid Modulated Monocytes via IL-6 and CCL2"

### Supplementary figures 1-8

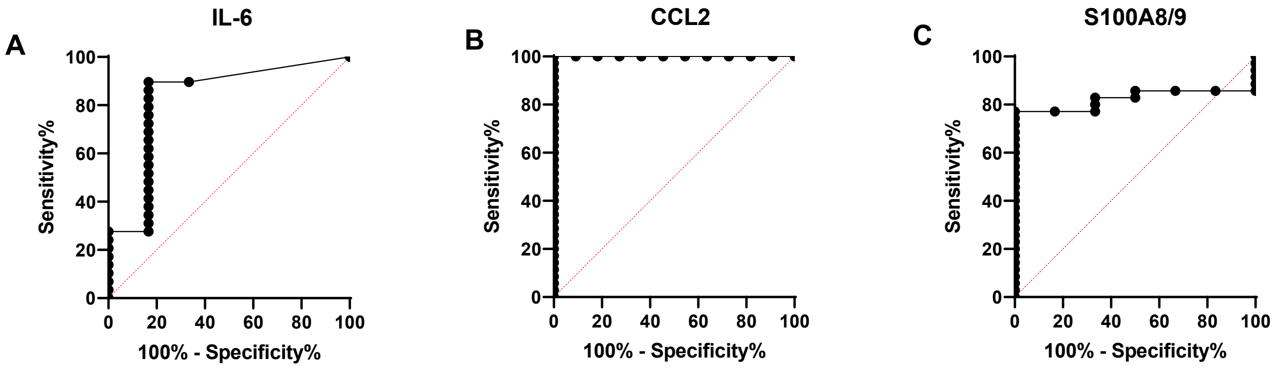


**Figure S1. IL-6, CCL2, and S100A8/9 are specifically increased in late OA SF, as indicated by the increased area under the receiver operating characteristic (ROC) curves**
**(A-C)** ROC curves indicated the specificity of IL-6, chemokine (C-C motif) ligand 2 (CCL2), and S100A8/9 measurements in late-OA synovial fluid (SF) compared to control, non-diseased SF samples; diagonal represents a random classifier.

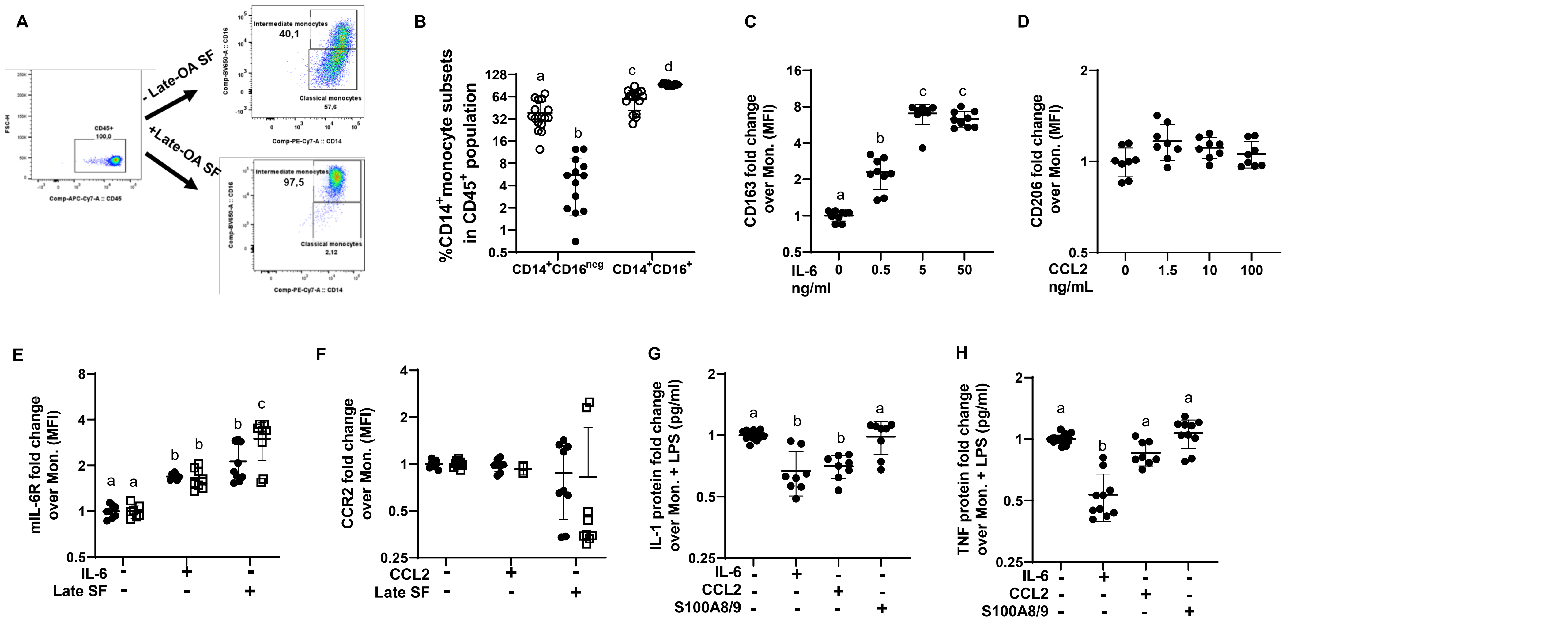


**Figure S2. Late-OA SF, IL-6, CCL2 and S100A8/9 effects on peripheral CD14⁺ monocytes**

**(A, B)** Frequency of classical (CD14⁺CD16^neg^) and intermediate (CD14⁺CD16⁺) peripheral healthy CD14⁺ monocyte subsets with (●) and without (○) OA synovial fluid (SF). Dose-response of healthy, peripheral CD14⁺ monocytes on **(C)** CD163 MFI with different IL-6 concentrations or on **(D)** CD206 MFI with different CCL2 concentrations. (**E, F**) membrane IL-6R (mIL-6R) and CCR2 expression on healthy (●) or KOA (□) CD14⁺ monocytes at indicated treatments. **(G, H)** TNF and IL-1 soluble factor production by CD14⁺ monocytes at indicated treatments, normalized to control LPS stimulation. N=3 biological replicates; n=3 technical replicates. Letters indicate significant differences based on an ordinary one-way ANOVA followed by Tukey's multiple comparisons test.

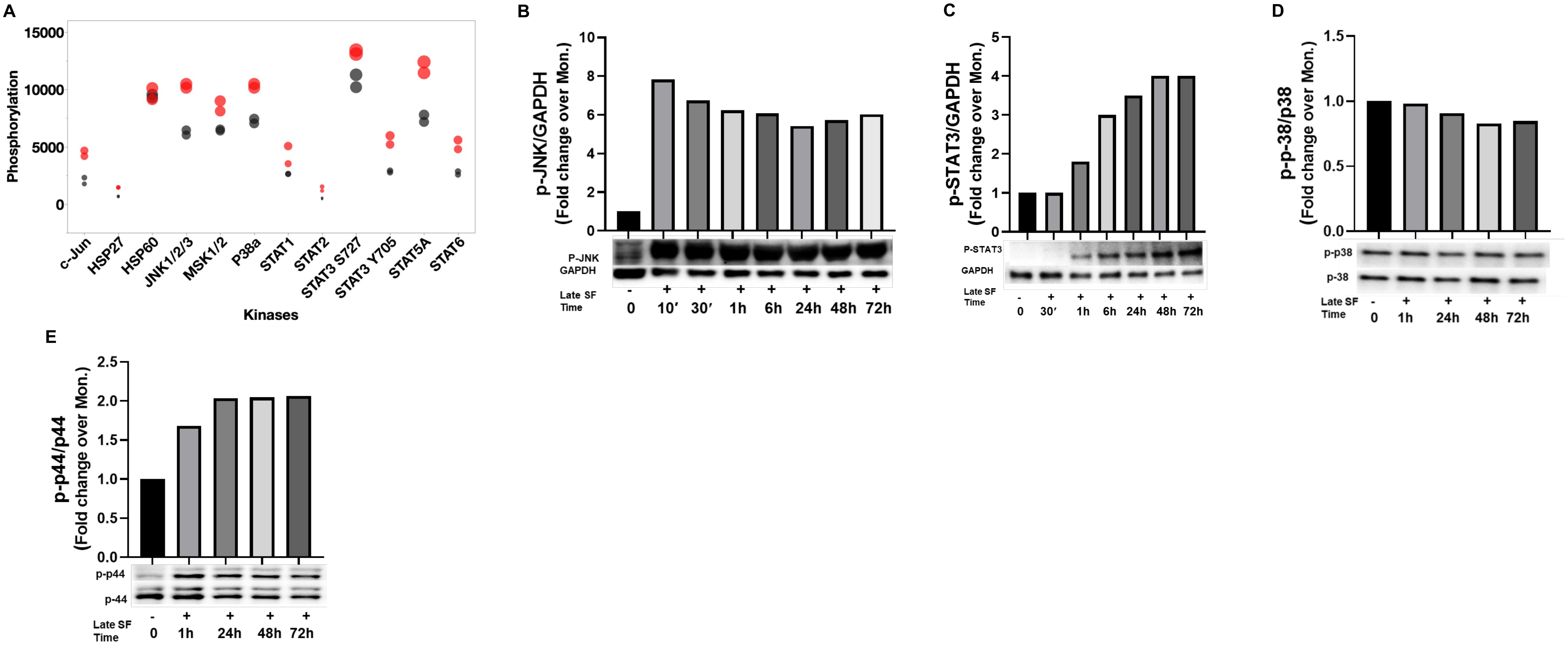


**Figure S3. Late-OA SF activates different signaling pathways in CD14⁺ monocytes**

**(A)** Bubble plot of the phosphorylation profiles of kinases in CD14⁺ monocytes treated with (red) and without late-OA SF (gray) with bubble size representing signal intensity corresponding to the level of phosphorylated kinase protein (N=1 biological replicates; n=2 technical replicates; small (<5000 pixels); medium (5000-10000 pixels); large (10000- 15000 pixels)). **(B-E)** Representative Western blots and bar graphs for phosphorylated (p)-JNK, p-STAT3 levels (normalized to GAPDH), and p-protein (p)38 levels (normalized to p38), p-p44 levels (normalized to p44) at the indicated times (N=1 donor, n=3 technical replicates). HSP27; Heat shock protein 27, HSP60; Heat shock protein 60, MSK 1/2; Mitogen and stress-activated kinases 1/2.

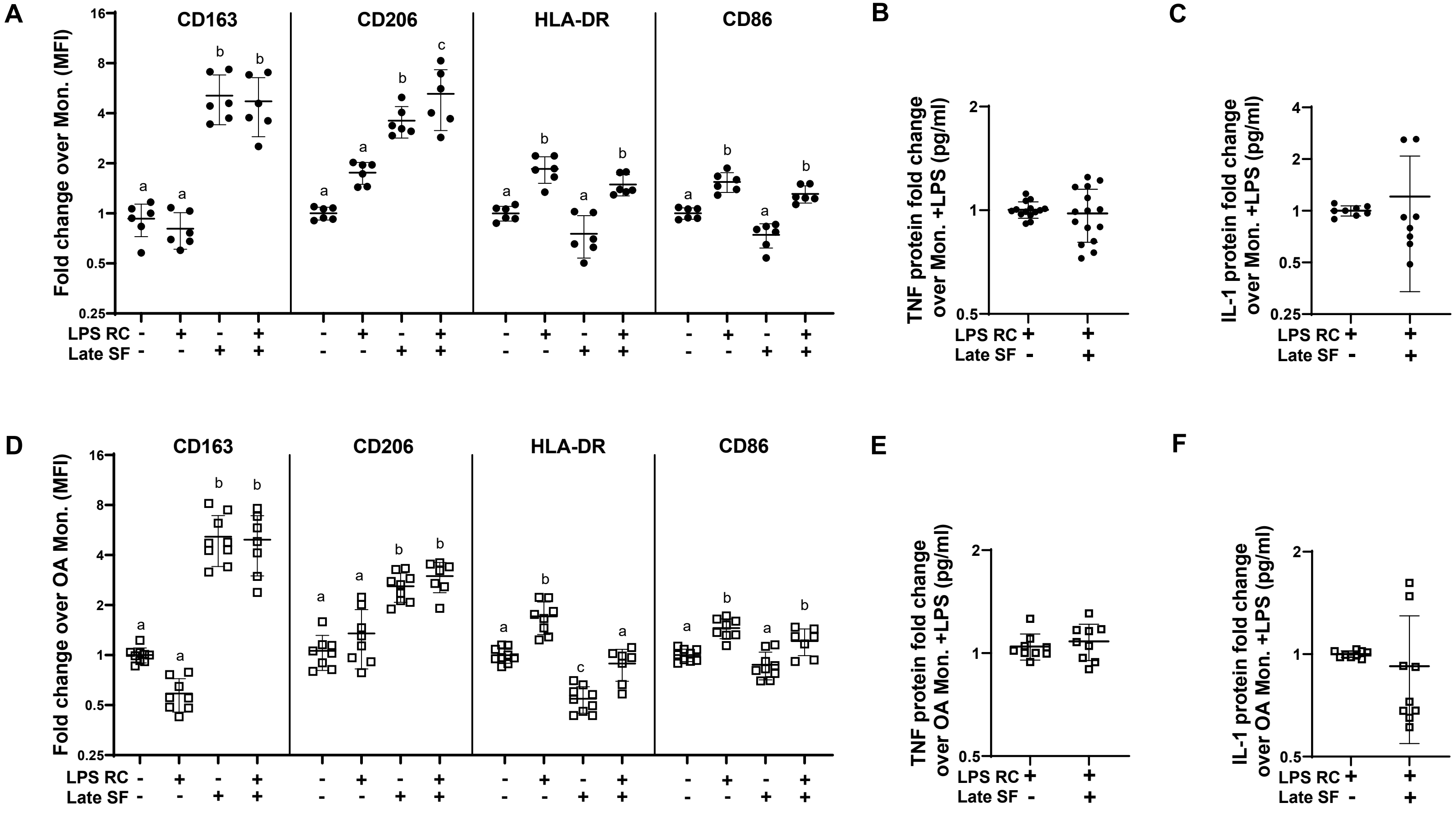


**Figure S4: CD14⁺ monocytes rechallenged with LPS in late-OA SF**

**(A, D)** CD163, CD206, HLA-DR, and CD86 MFI in healthy (●) or KOA (□) CD14⁺ monocytes exposed to pooled late OA SF after LPS re-challenge (RC)**. (B, E)** TNF, and **(C, F)** IL-1 soluble factors production from healthy (●) or KOA (□) peripheral CD14⁺ monocytes after LPS RC in late-OA SF. N=3 biological replicates; n=3 technical replicates. Letters indicate significant differences based on an ordinary one-way ANOVA followed by Tukey’s multiple comparisons test.

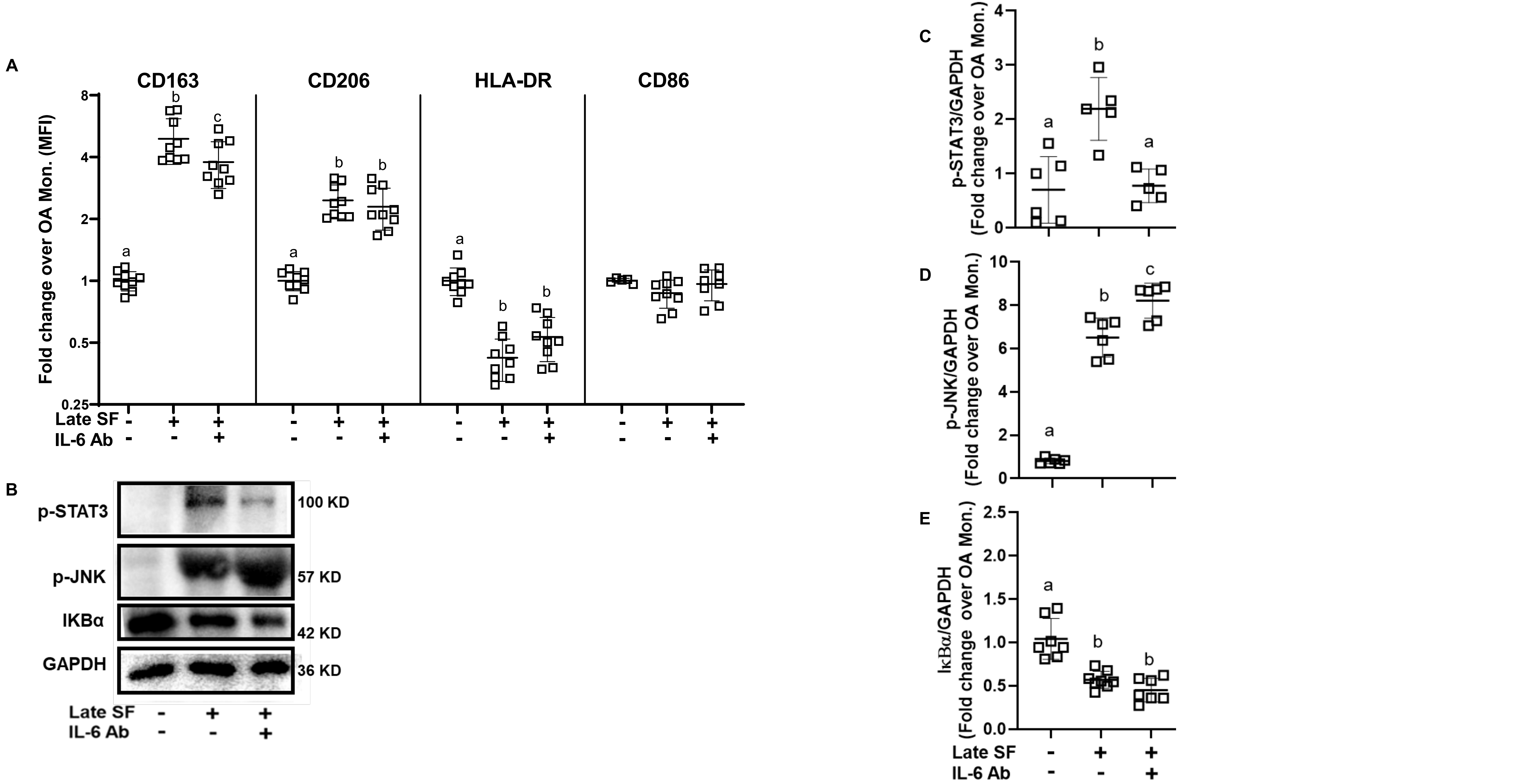


**Figure S5. Late-OA synovial fluid effects on KOA patient-sourced peripheral CD14⁺ monocytes**

**(A)** MFI of CD163, CD206, HLA-DR, and CD86 on KOA peripheral $CD14^+$ monocytes at indicated treatments. **(B)** Representative Western blot. **(C-E)** Bar graphs of p-STAT3, p-JNK, and IκBα levels (normalized to GAPDH). N=3 biological replicates; n=3 technical replicates. Letters indicate significant differences based on an ordinary one-way ANOVA followed by Tukey's multiple comparisons test.

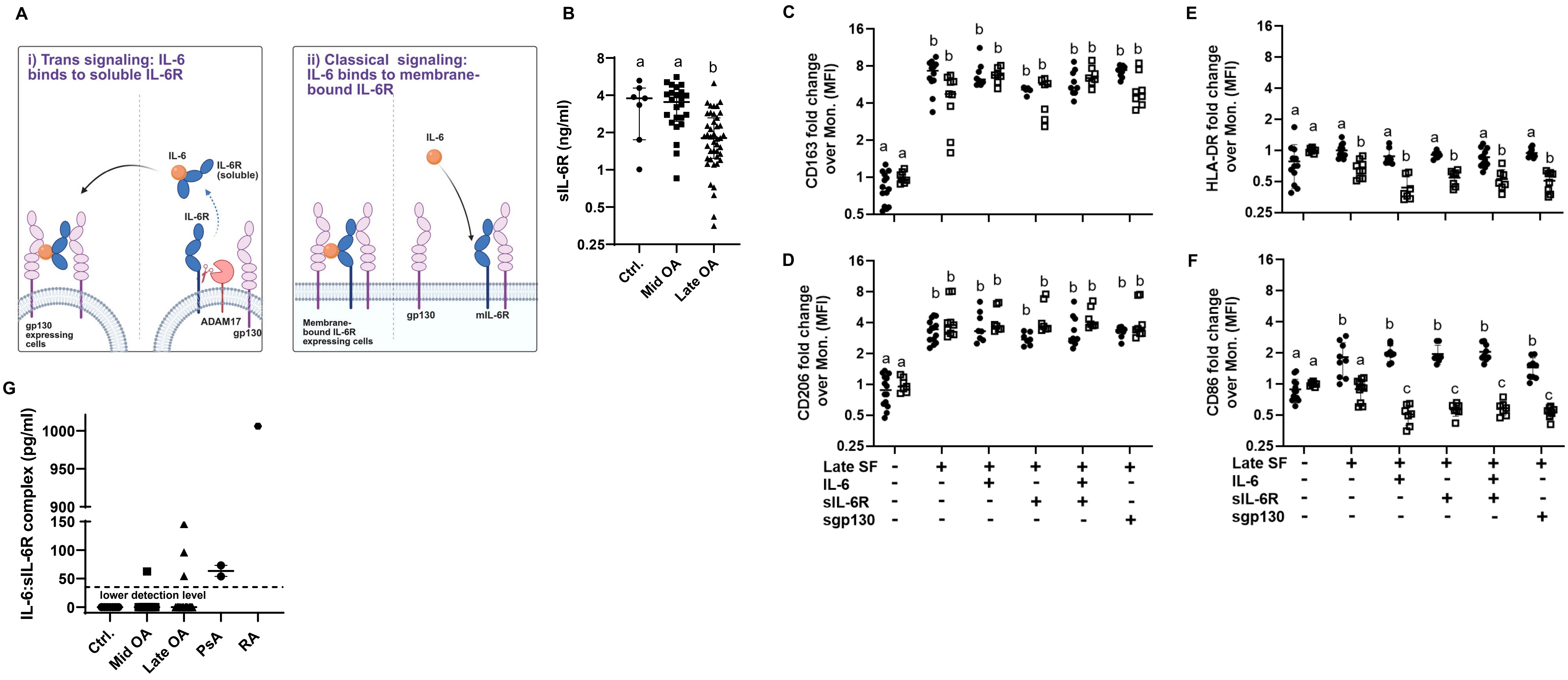


**Figure S6. Physiological levels of IL-6 in KOA SF signal through classical, not trans-signaling**

**(A)** IL-6 signaling in monocytes/MΦs: i) trans-signaling through glycoprotein (gp)130 and soluble IL-6R (sIL-6R shed by a disintegrin and metalloproteinase 17 (ADAM 17)); ii) classical signaling through membrane-bound (m)IL-6R and gp130. **(B)** Median and interquartile range of sIL-6R concentration in SF from ctrl. (non-diseased, N=7); mid-OA (N=24); and late-OA (N=43). **(C-F)** MFI expression of CD163, CD206, HLA-DR, and CD86 at indicated treatments in healthy (●) and KOA (□) $CD14^+$ monocytes; N=3 biological replicates; n = 3 technical replicates. **(G)** IL-6:sIL-6R complex levels in ctrl. (non-diseased; N=7); mid-OA (N=24); late-OA (N=72), psoriatic arthritis (PsA) SF (N=2) and rheumatoid arthritis (RA) SF (N=1). Letters indicate significant differences based on an ordinary one-way ANOVA followed by Tukey's multiple comparisons test.

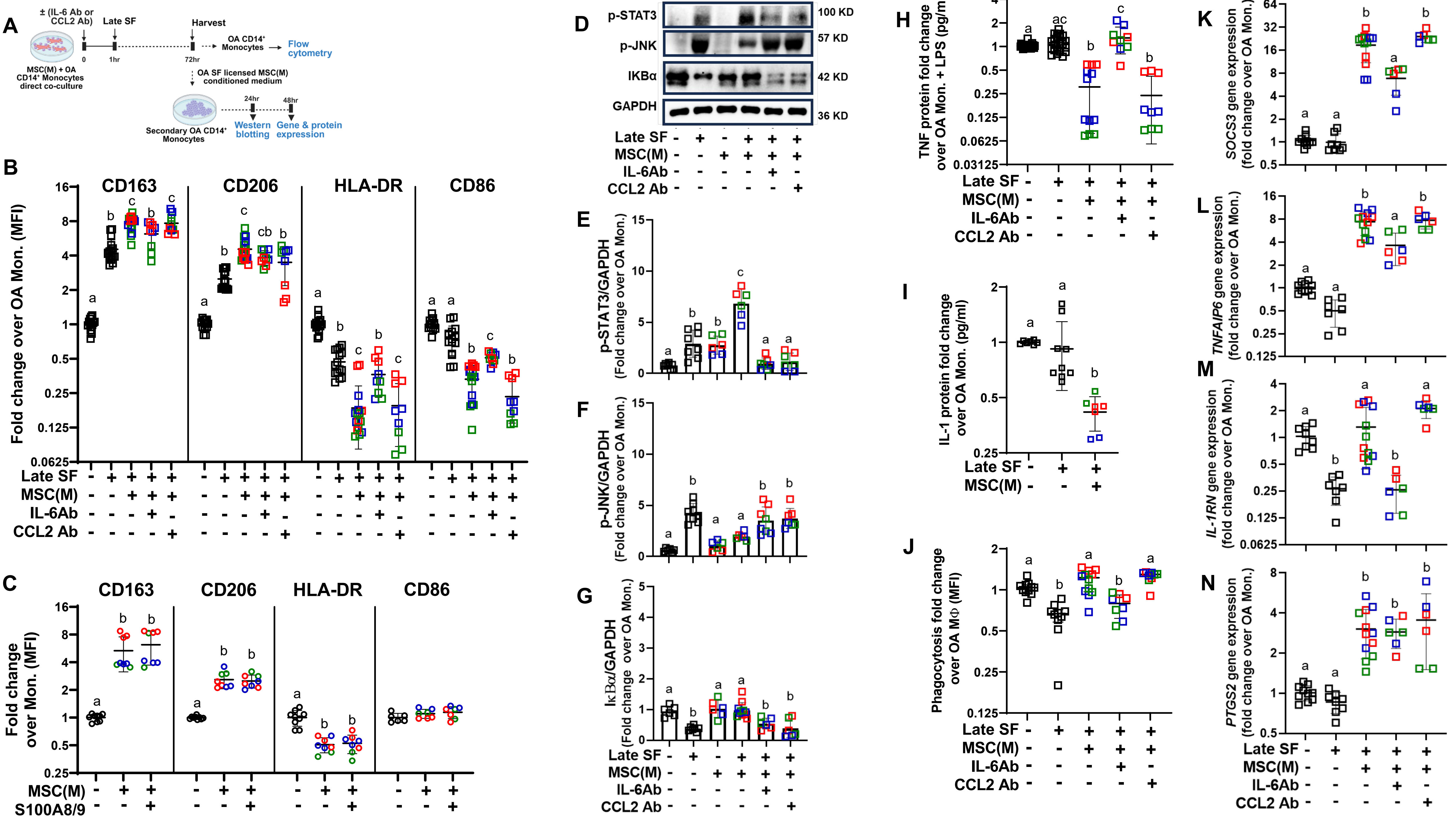

A
± (IL-6 Ab or CCL2 Ab)
Late SF
Harvest
MSC(M) + OA CD14+ Monocytes direct co-culture
0
1hr
72hr
OA CD14+ Monocytes
Flow cytometry
OA SF licensed MSC(M) conditioned medium
24hr
48hr
Secondary OA CD14+ Monocytes
Western blotting
Gene & protein expression
B
CD163
CD206
HLA-DR
CD86
Fold change over OA Mon. (MFI)
Late SF
MSC(M)
IL-6Ab
CCL2 Ab
C
Fold change over Mon. (MFI)
MSC(M)
S100A8/9
D
p-STAT3
p-JNK
IKBα
GAPDH
100 KD
57 KD
42 KD
36 KD
E
p-STAT3/GAPDH (Fold change over OA Mon.)
F
p-JNK/GAPDH (Fold change over OA Mon.)
G
IκBα/GAPDH (Fold change over OA Mon.)
H
TNF protein fold change over OA Mon. + LPS (pg/ml)
I
IL-1 protein fold change over OA Mon. (pg/ml)
J
Phagocytosis fold change over OA MΦ (MFI)
K
SOCS3 gene expression (fold change over OA Mon.)
L
TNFAIP6 gene expression (fold change over OA Mon.)
M
IL-1RN gene expression (fold change over OA Mon.)
N
PTGS2 gene expression (fold change over OA Mon.)

**Figure S7. Direct co-culture of MSC(M) with KOA CD14$^+$ monocytes in late-OA SF**

**(A)** Experimental schematic of KOA CD14$^+$ monocytes directly co-cultured with MSC(M); late-OA SF or IL-6/CCL2 neutralizing antibodies. **(B)** MFI of CD163, CD206, HLA-DR, and CD86, with or without late-OA SF. **(C)** CD163, CD206, HLA-DR, and CD86 MFI of healthy CD14$^+$ monocytes co-cultured with MSC(M) in the presence of S100A8/9. **(D)** Representative Western blot. **(E-G)** p-STAT3, p-JNK, and IκBα levels (normalized to GAPDH ). **(H, I)** Normalized TNF and IL-1 soluble factors production at indicated treatments. **(J)** Phagocytosis, at indicated treatments. **(K-N)** Expression of *SOCS3*, *TNFAIP6*, *IL-1RN, PTGS2 genes* . MSC(M) donors are represented by different colors (blue, green, red). N=3 biological replicates; n=3 technical replicates. Letters indicate significant differences based on an ordinary one-way ANOVA followed by Tukey's multiple comparisons test.

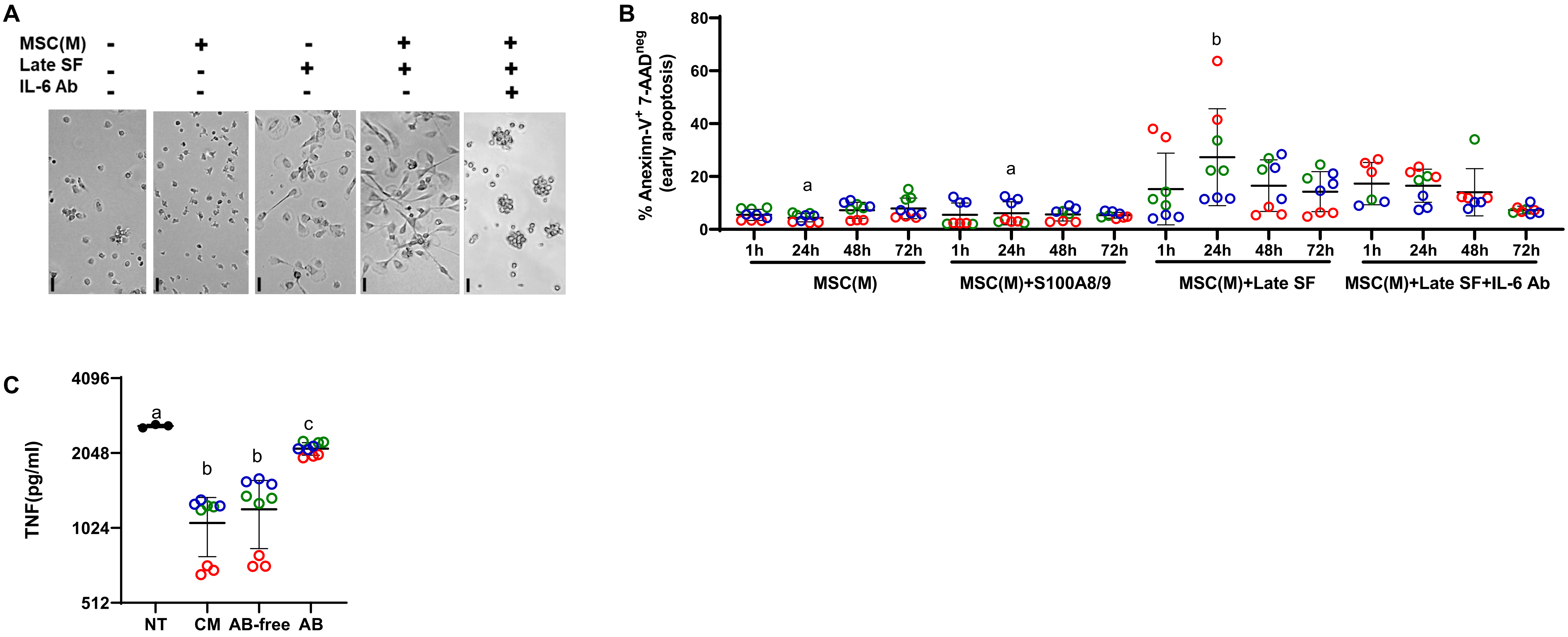


**Figure S8. Effects of MSC(M) soluble factors on secondary CD14+ monocytes**

**(A)** Representative CD14+ monocyte images at indicated treatments; 20x objective; scale bar, 10 µm. **(B)** Apoptosis frequency in MSC(M) treated with S100A8/9, late-OA SF, and late-OA LSF with IL-6-neutralizing antibody (Ab) at different time points. **(C)** TNF soluble factor production in response to different fractions of the licensed MSC(M) conditioned medium (CM), NT (untreated control), AB (apoptotic bodies), and AB-free CM. MSC(M) donors are represented by different colors (blue, green, red). N=3 biological replicates; n=3 technical replicates. Letters indicate significant differences based on an ordinary one-way ANOVA followed by Tukey's multiple comparisons test.

**Supplementary Tables**

**Tables S1-S9**

**Table S1. Demographics of KOA patients SF donors MSC(M) donors**

| **Variable** | **Mid OA patients (N = 35)** | **Late OA patients (N = 104)** | **MSC(M) donors (N=3)** |
|---|---|---|---|
| **BMI, Median (IQR)** | 29.51 (25.7-32.3) kg/$m^2$ | 29.34 (2.32 - 33.88) kg/$m^2$ | 25.2, 28, 29 kg/$m^2$ |
| **Age, Median (IQR)** | 57 (52 - 65) years | 65.5 (61 - 72) years | 32, 50, 58 years |
| **Male N, %** | 18 (51%) | 44 (42%) | 3 (100%) |
| **KL** | II, Early III | Late III, IV | Healthy |

**Table S2. IL-6, CCL2, and S100A8/9 measurements in SF**

| **IL-6 pg/mL** | | | **CCL2 pg/mL** | | | **S100 A8/9 ng/mL** | | |
|---|---|---|---|---|---|---|---|---|
| **control** N=6 | **Mid OA** N=21 | **Late OA** N=29 | **control** N=11 | **Mid OA** N=27 | **Late OA** N=35 | **control** N=6 | **Mid OA** N=30 | **Late OA** N=35 |
| 23.12 | 0 | 0 | 30.05 | 652.2 | 354.8 | 240.77 | 101.50 | 1762.60 |
| 0 | 33.99 | 540.09 | 26.37 | 597.6 | 934.6 | 179.11 | 195.60 | 1564.99 |
| 0 | 34.19 | 611.74 | 120.84 | 606.9 | 372.4 | 184.45 | 122.64 | 207.29 |
| 439.64 | 73.6 | 690.3 | 103.07 | 312.9 | 723.7 | 187.34 | 190.30 | 940.49 |
| 0 | 75.435 | 151.11 | 62.6 | 347.4 | 701.4 | 256.81 | 74.41 | 272.42 |
| 0 | 275.24 | 88.34 | 70.7 | 1112.4 | 1392.4 | 205.22 | 110.27 | 387.76 |
| | 0 | 67.14 | 28.11 | 246 | 548.9 | | 160.07 | 1368.60 |
| | 119.70 | 93.95 | 23.77 | 464.5 | 332.1 | | 291.71 | 1178.33 |
| | 0 | 232.06 | 119.59 | 335.8 | 412.1 | | 127.49 | 1111.85 |
| | 67.37 | 306.06 | 79.59 | 255.6 | 208.3 | | 193.56 | 597.42 |
| | 37.29 | 481.65 | 74 | 516.3 | 447.8 | | 861.93 | 942.93 |
| | 293.55 | 108.66 | | 559.4 | 1248.1 | | 1142.32 | 116.61 |
| | 730.93 | 150.71 | | 305.8 | 279 | | 181.64 | 272.69 |
| | 0 | 327.90 | | 813.4 | 287.4 | | 160.77 | 435.72 |
| | 688.88 | 233.10 | | 181.2 | 846.1 | | 111.24 | 1035.43 |
| | 0 | 0 | | 58.9 | 268.7 | | 223.6 | 261.86 |
| | 449.21 | 338.95 | | 112.2 | 1353.4 | | 112.79 | 841.19 |

| | | | | | | | | |
|---|---|---|---|---|---|---|---|---|
| | 445.37 | 758.07 | | 1007.1 | 451.4 | | 26.62 | 160.90 |
| | 40.2 | 76.28 | | 162.9 | 812.6 | | 360.68 | 666.70 |
| | 0 | 60.74 | | 101.6 | 159.3 | | 69.401 | 427.80 |
| | | 51.015 | | 216.4 | 299.9 | | 134.71 | 348.71 |
| | | 31.09 | | 329.9 | 243.9 | | 628.67 | 114.81 |
| | | 895.52 | | 597.3 | 427.8 | | 250.07 | 453.05 |
| | | 0 | | 268.1 | 983.7 | | 1003.08 | 193.06 |
| | | 584.85 | | 290.9 | 155.1 | | 771.81 | 105.03 |
| | | 752.935 | | 190.8 | 242 | | 544.68 | 205.94 |
| | | 297.65 | | 225.2 | 235.2 | | 167.42 | 1377.94 |
| | | 119.785 | | | 278.8 | | 384.44 | 325.29 |
| | | 175.65 | | | 475.1 | | 857.72 | 1756.22 |
| | | | | | 715.8 | | 142.71 | 1948.25 |
| | | | | | 435.7 | | | 372.39 |
| | | | | | 334.6 | | | 1080.34 |
| | | | | | 937.5 | | | 859.42 |
| | | | | | 122.9 | | | 493.54 |
| | | | | | 338.9 | | | 105.7 |

**Table S3. CD14⁺ monocytes from healthy or KOA patients in response to late OA SF**

| Measure | Healthy CD14⁺ Monocytes + Late-OA SF rel. to untreated controls | OA CD14⁺ Monocytes +Late-OA SF rel. to untreated controls |
|---|---|---|
| **CD163 MFI** | ↑ | ↑ |
| **CD206 MFI** | ↑ | ↑ |
| **HLADR MFI** | ↔ | ↓ |
| **CD86 MFI** | ↔ | ↔ |
| **mIL-6R MFI** | ↑ | ↑ |
| **CCR2 MFI** | ↔ | ↔ |
| **pSTAT3/GAPDH** | ↑ | ↑ |

| | | |
|---|---|---|
| **pJNK/GAPDH** | ↑ | ↑ |
| **IKB/GAPDH** | ↓ | ↓ |
| ***SOCS3*** | ↔ | ↔ |
| ***TNFAIP6*** | ↔ | ↔ |
| ***IL1RN*** | ↔ | ↓ |
| ***PTGS2*** | ↔ | ↔ |
| **TNF secretion** | ↔ | ↔ |
| **IL1 secretion** | ↔ | ↔ |
| **Phagocytosis** | ↓ | ↓ |

**Table S4: Analysis of CD14$^+$ monocytes with tested stimulators**

| | **Rel. to untreated CD14$^+$ Monocytes** | | | | | **Rel. to Late-OA SF treated CD14$^+$ Monocytes** | | |
|---|---|---|---|---|---|---|---|---|
| | **Late-OA SF** | **IL-6** | **CCL2** | **S100A8/9** | **IL-6 + CCL2 + S100A8/9** | **IL-6 Ab** | **CCL2 Ab** | **TAK242** |
| **CD206** | ++ | + | = | = | = | = | - | = |
| **CD163** | +++ | + | = | = | = | - | = | = |
| **HLA-DR** | = | = | = | = | = | = | = | = |
| **CD86** | = | = | = | = | = | = | = | = |
| **p-STAT3** | + | + | = | = | = | -- | - | = |
| **p-JNK** | ++ | = | = | = | = | + | = | = |
| **IκBα** | - | = | = | - | = | - | = | + |
| **TNF** | = | NA | NA | NA | - | + | = | - |
| **IL-1** | = | NA | NA | NA | = | + | = | = |
| **Phagocytosis** | - | NA | NA | NA | - | = | = | = |

**Table S5: Analysis of MSC(M) effects on CD14$^+$ monocytes**

| | **Rel. to Late-OA SF** | **Rel. to MSC + Late-OA SF treated cells** | |
|---|---|---|---|
| | **Effect of MSC + LSF** | **Effect of IL-6 Ab** | **Effect of CCL2 Ab** |
| **CD206** | +++ | = | - |
| **CD163** | ++++ | - | - |
| **HLA-DR** | - | + | = |

| | | | |
|---|---|---|---|
| **CD86** | - | = | = |
| **p-STAT3** | ++ | -- | -- |
| **p-JNK** | - | ++ | ++ |
| **IκBα** | + | - | - |
| **IL-1** | - | NA | NA |
| **TNF** | - | ++ | = |
| **Phagocytosis** | + | - | = |
| ***IL1RN*** | + | - | = |
| ***TNFAIP6*** | + | - | = |
| ***PTGS2*** | + | = | = |
| ***SOCS3*** | + | -- | - |

**Table S6. Antibodies for immunophenotyping $CD14^+$ peripheral monocytes and SF monocytes/MΦs**

| Antibodies | Source | Dilution | Catalogue number (Cat#) |
|---|---|---|---|
| APC-Cy7 anti-human CD45 | BioLegend | 1:100 | Clone #HI30; Cat# 304014 |
| PE-Cy7 anti-human CD14 | BioLegend | 1:100 | Clone #M5E2; Cat# 982510 |
| BV650 anti-human CD16 | BioLegend | 1:50 | Clone #3G8; Cat# 302042 |
| FITC anti-human CD163 | BioLegend | 1:50 | Clone #GHI/61; Cat# 333618 |
| BV421 anti-human CD206 | BioLegend | 1:50 | Clone #15-2; Cat# 321126 |
| PerCP-Cy5.5 anti-human HLA-DR | BioLegend | 1:100 | Clone #L243; Cat# 980414 |
| PerCP-Cy5.5 anti-human mIL6Ra | BioLegend | 1:50 | Clone #UV4; Cat# 352812 |
| PE anti-human TNF antibody | BioLegend | 1:50 | Clone #MAb11; Cat# 502909 |
| Pacific Blue anti-human IL-1 | BioLegend | 1:50 | Clone #H1b-98; Cat# 511710 |
| CCR2 | BioLegend | 1:50 | Clone #KO36C2; Cat# 357210 |

**Table S7. ELISA kits to detect soluble factors in SF and conditioned medium**

| Soluble factors | Source | Dilution Factor | Catalogue number |
|---|---|---|---|
| IL-6 | Biolegend | SF,1:5 | 430504 |
| CCL2 | Biolegend | SF,1:25 | 438804 |
| S100A8/9 | R&D Systems | SF,1:250 | DY8226-05 |
| IL6-sIL6R | R&D Systems | SF, not diluted | DY8139-05 |
| sIL-6R | My BioSource | SF,1:2 | MBS266072 |
| IL-1 | Biolegend | CM, not diluted | 437004 |
| TNFα | R&D Systems | CM,1:2 | DY210-05 |

**Table S8. Antibodies for immunoblotting**

| Antibodies | Source | Catalogue number |
|---|---|---|
| Phospho-STAT3-Y705 Rabbit mAb | ABclonal | AP0705 |
| Phospho-JNK1/2-T183/Y185 Rabbit mAb | ABclonal | AP0473 |
| Phospho-p38 MAP-T180/Y182 Rabbit mAb | ABclonal | AP0526 |
| P38 MAPK Rabbit mAb | Cell Signaling | 9212 |
| p-p44/42 (T202/Y204) (E10) mouse Ab | Cell Signaling | 9106 |
| P44/42 MAPK Erk1/2 (137F5) Rabbit mAb | Cell Signaling | 4695 |
| IKB (L35A5) mouse Ab | Cell Signaling | 4814 |
| GAPDH (14C10) Rabbit mAb | Cell Signaling | 2118 |

**Table S9. Primer sequences**

| Genes name | Forward sequence | Reverse sequence |
|---|---|---|
| *GAPDH* | GGAGCGAGATCCCTCCAAAAT | GGCTGTTGTCATACTTCTCATGG |
| *B2M* | CTCCGTGGCCTTAGCTGTG | TTTGGAGTACGCTGGATAGCCT |
| *β-Actin* | CTCACCATGGATGATGATATCGC | AGGAATCCTTCTGACCCATGC |
| *PTGS2* | ATAAGCGAGGGCCAGCTTTC | CGCAGTTTACGCTGTCTAGC |
| *SOCS3* | TTCGGGACCAGCCCCC | AACTTGCTGTGGGTGACCAT |
| *TNFAIP6* | AGCACGGTCTGGCAAATACA | ATCCATCCAGCAGCACAGAC |
| *IL-1RN* | AGC ATG AGG CTC AAT GGG TA | AAA TCC AGC AAG ATG CAA GC |
