## Supplementary figures and images for "Mesenchymal Stromal Cells Immunosuppress Osteoarthritis Synovial Fluid Modulated Monocytes via IL-6 and CCL2"

### Graphical Abstract

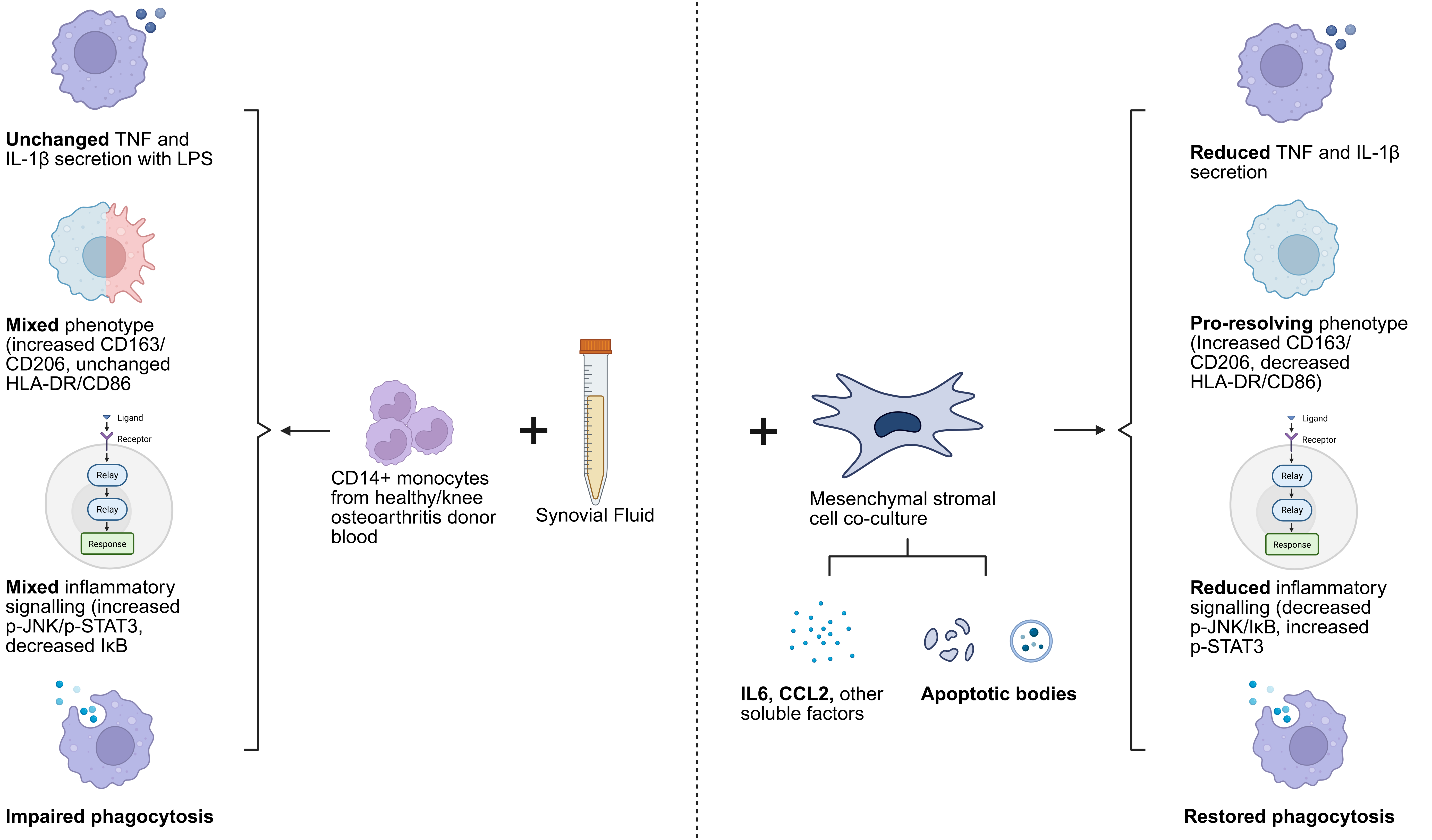
